# Trait-specific indirect effects underlie variation in the response of spiders to cannibalistic social partners

**DOI:** 10.1101/2022.12.08.519483

**Authors:** Jorge F. Henriques, Mariángeles Lacava, Celeste Guzman, Maria Pilar Gavin-Centol, Dolores Ruiz-Lupión, Alberto Ruiz, Carmen Viera, Jordi Moya-Laraño, Sara Magalhães

## Abstract

Organisms may respond in different ways to the risk posed by conspecifics, but the cause of such variation remains elusive. Here, we use a half-sib/full-sib design to evaluate the contribution of (indirect) genetic or environmental effects to the behavioral response of the cannibalistic wolf spider *Lycosa fasciiventris* (Dufour, 1835) towards conspecific cues. Spiders showed variation in relative occupancy time, activity, and velocity on patches with or without conspecific cues, but direct genetic variance was only found for occupancy time. These three traits were correlated and could be lumped in a principal component: spiders spending more time in patches with conspecific cues moved less and at a lower rate in those areas. Genetic and/or environmental components of carapace width and weight loss in the social partner were significantly correlated with the principal component of focal individuals. Variation in these traits may reflect the quality and/or quantity of cues produced by social partners, hence focal individuals were likely behaving along a continuum of strategies in response to the risk posed by social partners. Therefore, environmental and genetic trait variation in the social partners may be key to maintain trait diversity in focal individuals, even in the absence of direct genetic variation.

## Introduction

The presence of others is one of the most common environmental effects that organisms face (Bailey and Zuk 2012; Bleakley et al. 2013; Rudin et al. 2018). Social partners are a particular environment because they contain genes themselves and can thus evolve. Indirect genetic effects (IGE) occur when genetic variation in social partners affect traits in focal individuals (Moore et al. 1997). Additionally, social partners may affect the behaviour of focal individuals via indirect environmental effects (IEE) whenever their effect on others occurs through their environmental variance (Moore et al. 1997). Both IGE and IEE can be substantial in magnitude. For instance, in chipmunks IGE and IEE explained a high proportion of variation in traits such as exploration or body mass in focal individuals (Santostefano et al. 2021).

IGE have the potential to affect the evolutionary trajectory of populations, as they may hamper or foster trait evolution (Moore et al. 1997; McGlothlin et al. 2010). For example, IGE mediate sexual conflict over egg laying time in gulls (Brommer and Rattiste 2008), potentially hampering trait change in these birds. In contrast, IGE and direct genetic effects (DGE) were positively correlated for this trait in song sparrows, which is expected to lead to evolution being faster due to IGE (Germain et al. 2016). IGE need to be stronger than DGE to affect evolutionary trajectories (Moore et al. 1997). Although this is not the case in several systems (Chakrabarty et al. 2019; Moiron et al. 2020; Ribeiro et al. 2020) it is still important to estimate these effects, because they may affect trait variation in focal individuals.

An important role of indirect effects is their contribution to the maintenance of diversity (Day and Bonduriansky 2011; Bailey et al. 2018). Indeed, it has been argued that interactions with other individuals may underlie the large values of phenotypic variation in communities (Shuster et al. 2006) of additive genetic variance for fitness in wild populations (Bonnet et al. 2022) as well as species coexistence in food webs (Moya-Laraño 2011; Whitlock et al. 2011). Whereas social partners can only affect trait evolution in other individuals via IGE, their impact on trait diversity can be mediated by IGE, IEE or anything in between (i.e., non-additive genetic effects, genetic or environmental maternal effects).

The canonical way by which IGE contributes to maintaining trait genetic diversity is whenever IGE and DGE have opposite effects on fitness, which occurs in cases of conflict. Indeed, IGE underlying sexual conflict may explain trait diversity (Marie-Orleach et al. 2017) and even resolve the lek paradox (Pischedda and Chippindale 2006; Miller and Moore 2007; Danielson-François et al. 2009). Moreover, IGE are predicted to maintain trait diversity in cases of parent-offspring conflict (Kölliker et al. 2005).

Trait diversity may also be maintained by social partners via more subtle mechanisms. For example, some genotypes are sensitive to IGE whereas others are not (Han et al. 2018; Carter et al. 2019). IGE may also trigger different responses in different genotypes. For example, variation in aboveground and belowground traits in plants is explained by the genetic composition of their neighbours (Genung et al. 2012). Also, mate choice may be affected by the genotypic composition of social partners in treehoppers (Rebar and Rodríguez 2013). Different levels of social interactions may also trigger different responses. For example, variation in levels of parental care and sibling competition affect variation in body size in burying beetles (Schrader et al. 2018). Conversely, mother body size exerts an indirect effect on brood size in dung beetles (Hunt and Simmons 2002). Moreover, the genetic composition of neighbours affects the expression of fighting characters in sea anemones (Lane et al. 2020) and variation in the exploratory behaviour of social partners explains variation in aggressiveness in Mediterranean field crickets (Santostefano et al. 2017*a*).

IGE and IEE have been particularly studied in the context of behavioural interactions. Indeed, social partners modulate several behaviours in conspecifics, such as boldness and exploratory behaviour (Rudin et al. 2018; Blake et al. 2018), mating behaviour (Marie-Orleach et al. 2017), aggressiveness (Wilson et al. 2009; Bleakley et al. 2013; Santostefano et al. 2017*b*; Lane et al. 2020) or the avoidance of predation or (kin) competition (Bleakley and Brodie IV 2009; Costa e Silva et al. 2013; Rode et al. 2017; Dewan et al. 2019).

Conspecific cues may be considered a trait by itself, as they have been recently demonstrated to be a source of IGE (Rode et al. 2017; Dewan et al. 2019). Thus, in principle, traits associated to the quantity and/or quality of the cues released could also be linked to IGE or IEE. Excreta are often used as cues by animals to avoid patches of relatively high predation risk (Persons and Rypstra 2001; Mogali et al. 2020). Candidate traits explaining the quantity or quality of excreta are morphological, such as body size, since larger animals release more excreta (Milanovich and Hopton 2016), or physiological, such as weight loss, which may be associated to higher metabolic rates, but also to lower assimilation efficiencies. In the context of cannibalistic interactions, conspecifics can use this information to avoid competition or predation by a cannibalistic competitor, or to evaluate the chances of an attack being successful. Therefore, assessing the role of IGE or IEE in the response to cues left by social partners (or to traits associated to their release of cues), may be paramount to understanding the prevalence and maintenance of different responses to conspecific cannibals.

Wolf spiders (Lycosidae) are cannibalistic generalist predators (Nentwig 1986) that follow a sit-and-pursue foraging strategy, by which they sit on patches waiting for their prey but they also often move between such patches (Schmitz and Suttle 2001). Upon a conspecific encounter during these foraging bouts, cannibalism usually depends on body size differences (Hvam et al. 2005; Rypstra and Samu 2005, 2006; Gonzalez 2012; Wise 2006). These spiders release silk and excreta (Persons and Rypstra 2001), which can be used as cues by conspecifics (Roberts and Uetz 2004). In cannibalistic species, wolf spider prey may respond in an anti-predatory fashion to the quantity (Persons et al. 2001) and quality (Persons and Rypstra 2000) of these cues. The variety of defensive tactics include reduced movement or velocity in patches with predator cues, or reduced occupancy of those patches (Persons et al. 2001). Even though wolf spiders are a model system to study cannibalism (Wise 2006), their response to the risk posed by conspecific cues has only been considered in a handful of studies, finding either no response (Barnes et al. 2002), that spiders are attracted to such cues (Wetter et al. 2012), or that they avoid areas with high conspecifics numbers (Hanna and Eason 2013). Such variation could be due to (a) cannibalism involves feeding and being fed upon, so different strategies may be adopted depending on whether individuals are predator or prey in that specific encounter and/or (b) different avoidance strategies may occur, from avoiding patches altogether to avoiding being seen in those patches. Moreover, the amount of food is known to modulate cannibalism in spiders (Wise 2006). Indeed, on the one hand, hungry spiders may be inclined to enter patches with conspecific cues to attack and feed upon conspecifics. On the other hand, patches with cues of well-fed spiders may be entered by other spiders because they are perceived as safe.

Here, we measure the impact of DGEs and/or IGEs on the behaviour of cannibalistic wolf spiders exposed to conspecific cues. Some spider traits have been shown to exhibit significant additive genetic variance (Bonte and Lens 2007; Hendrickx et al. 2008), but genetic and non-genetic maternal effects also explain large parts of trait variance (Bonte et al. 2006, 2007; Bonte and Lens 2007; Hendrickx et al. 2008; Storm and Lima 2010; Shannon et al. 2022). To account for this, we use a paternal half-sib/full-sib split-brood design to measure occupancy time, activity, and velocity on patches with or without conspecific cues in the *Lycosa fasciiventris* (Dufour, 1835), a non-burrowing wolf spider with an annual life-cycle, inhabiting semi-arid lands in the Iberian Peninsula (Barrientos 2004; Planas et al. 2013; Gavín-Centol et al. 2017), which shows size-dependent cannibalism (Henriques 2020). We then tested whether behavioural traits in focal individuals were affected by DGE, DEE or IGE and IEE from social partners. We hypothesize that focal individuals will avoid patches with conspecifics, and that the magnitude of this response will be affected by the genetic or environmental component of traits associated with cue emission in social partners.

## Material and Methods

### Spider collection and rearing conditions

*Lycosa fasciiventris* individuals (adult males and subadult females) were collected between June and July 2015 in four different localities within the Almeria province (South-East Spain), in dry temporal washes (“ramblas”): near Boca de los Frailes (36.8036°N, 2.1386°O), near Carboneras town (36.9667°N, 2.1019°O), at Almanzora river (37.3414°N, 2.0078°O) and near Paraje las Palmerillas, Estación Experimental Cajamar (36.7917°N, 2.6891°O). They were reared in individual tanks (22 cm x 18 cm x 18 cm) with the bottom filled with 2-3 cm of soil collected from one of the sites. All individuals were fed once a week with size-matched crickets (*Gryllus assimilis* Fabricius, 1775) purchased from the pet supply virtual store Exofauna (available at: https://exofauna.com). Spiders had access to water *ad libitum* through a 40 ml vial filled with water and covered with cotton. Vials were checked and refilled every 2-3 days. Holding tanks were placed in a climate chamber that simulated outdoor climatic conditions of the preceding weekly average conditions in the Almeria province (night:day temperature cycles between 18.7 - 34.3 ºC; photoperiod of 17:7 h - 16:8 h light-dark with light bulbs of 54W and a relative humidity of 50 -65 %).

Individuals used in the experiments were removed from their mothers’ back 42 ± 8 days after hatching and placed in separate cylindrical containers (15 cm height and 6 cm of diameter) inside a growth chamber with controlled temperature (25°C ± 1ºC), humidity (70% ± 5%) and photoperiod (16:8 hours light:dark). The bottom of each container was covered with a filter paper replaced weekly. Water was provided *ad libitum* from a cotton string submerged in a reservoir, below each container and providing water by capillarity (Moskalik and Uetz 2011). Each week, spiderlings were fed with fruit flies (*Drosophila melanogaster* Meigen, 1830), reared in a nitrogen rich medium supplemented with high quality dogfood to ensure survival and growth of spiderlings (Jensen et al. 2011). A portion of the offspring within each dam family (3 out of 12) was reared in a richer environment by providing them three times the amount of food given to the remaining spiders, to examine how this affected both cue emission and the response to such cues. For logistic reasons, spiders were not reared until maturation and thus their sex was unknown. We assumed that sex differences were not very pronounced at the early stages, as it is the case in a syntopic and co-generic wolf spider (e.g., Fernández-Montraveta and Moya-Laraño 2007).

### Experiment

We aimed at testing the role of genetic and indirect genetic effects in shaping the variability in the interaction between cue-emitting spiders and spiders that perceive such cues. Our experimental design is thus highly asymmetrical, as one individual is only present via the cues it emits, whereas behaviour is measured in the other individual. Note also that each individual served first as a social partner, emitting cues, and was subsequently used to measure its behaviour in another arena. All trials were recorded between April 21^st^ and June 8^th^ 2016 in a total of 50 blocks and 17 recording days. Blocks consisted of 3 rows x 5 columns = 15 Petri dishes (individuals) recorded under a single camera. Usually, three blocks were recorded in a single day with three different cameras. More details are provided below.

#### Cues and traits in social partners

Behavioral responses to conspecific cues were measured in small Petri dishes (5.5 cm diameter) with the bottom covered with filter paper divided in two halves: one containing intact filter paper (control) and the other impregnated with conspecific cues (e.g., excreta and silk). These cues were produced by juvenile conspecifics (hereafter social partners) enclosed in a small Petri dish for 10 days with filter paper on the bottom and fed with 10 fruit flies each during the first 36 hours. The nature of the cues released included the excreta, silk, odour and chemo-tactile cues of dead prey. The spatial position of the two filter paper halves was randomized to eliminate any potential side bias.

As a proxy for the amount of cues produced, we used (a) weight loss of spiders confined within a petri dish for 36h and (b) carapace width, reflecting body size. In spiders, weight loss may be an indication of the quantity or quality of conspecific cues released. Indeed, since a great proportion of the cues likely correspond to excreta, animals losing more weight were likely those that also released more cues. Alternatively, higher weight loss may be related to animals that have higher voracities, as recently found in another wolf spider (Rádai et al. 2017), which could be associated to their willingness to attack conspecifics (Arnqvist and Henriksson 1997). Moreover, carapace width is known to be correlated with the amount of cues produced in spiders (Persons et al. 2001).

Weight loss was calculated by feeding spiders with 10 flies each, then weighting them after 36 hours and after 10 days. Proportional weight loss was then estimated as the difference in weight between these two measures divided by the initial weight. During this period, spiders were provided water *ad libitum* by means of a soaked piece of cotton of about 1 cm in diameter, which was re-soaked every two days. Body mass was measured to the nearest 0.1 mg using a high precision scale (Mettler Toledo XP26). Body size was assessed by measuring the maximum carapace width (i.e., carapace width at its maximum span). Measurements were performed with a stereomicroscope (Leica MZ125) with a precision of 0.1 mm.

#### Behavioral measurements

Spiders in which the behavioral response to social cues was measured (hereafter focal individuals) were randomly assigned to a Petri dish, but care was taken that they were not genetically related to and came from the same feeding treatment as their social partner. Note that each individual was used both as social partner and focal individual.

During the behavioral observations, light was made homogeneous by introducing the camera and the Petri dishes inside a 40×40×30 styrofoam box. Locomotor behavior was measured by monitoring spiders, in blocks of 15, through recordings retrieved from a video camera (Sony® HDR CX-150) placed overhead. A video-tracking software was implemented, allowing estimating movement at 25 frames/second. We obtained information for 563 spiders, with a mode recording time of 6 hours for each spider (min = 0.38 h, max = 6.04 h, median = 5.5 h). This large variation in recording time was due to an ant or fly walking on top of the Petri dish, or to a lack of space in the camera hard drive. However, we included recording time in the statistical analyses (see below), and it did not affect our results.

Differences in behavioral patterns between sides were estimated using the relative interaction intensity (RII) index described in Armas et al. (2004):

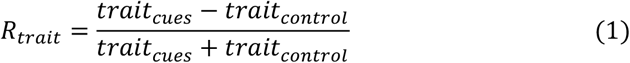

where *trait*_*cues*_ is the mean value of the trait in the patch with cues and *trait*_*control*_ is the mean value of the trait in the patch without cues. Using this general formula, we calculated: (a) the relative activity (RA) index as the time spent moving, (b) the relative mean velocity (RMV) index as the difference in speed and (c) the relative occupancy time (ROT) index as the difference in time spent in patches with or without conspecific cues (fig. S1).

In addition to these three traits, we performed a principal component analysis (PCA) to estimate a composite behavioural score based on the three traits, thus accounting for potential correlations among them. This was done in R v.4.0.2 (R Core Team 2020), using the function “principal” in library “psych” without axes rotation (fig. S2). We sought for a single PC that explained > 50% of the variance and in which the three traits had substantial loadings (, ≥ |0.7|).

In these focal individuals, we also used the measurement of weight loss and carapace width, as described above (cf. Cues and traits in social partners). Moreover, we added a measure of body condition, i.e., the maximum abdomen width (where nutrients and body fats are stored). Body condition has been shown to be negatively associated to movement in these spiders (Moya-Larano 2002), hence we used them as covariates in the statistical models measuring DGE, such as to account for this potential source of variation (see below).

### Quantitative genetic estimations

Both focal individuals and social partners come from the same paternal half-sib/full-sib breeding design (Lynch and Walsh 1998; Falconer and Mackay 1996), in which 52 males were each mated with two virgin (N=104) females to generate paternal half-sib families. This allowed us to calculate DGE and DEE on traits involved in the behavioral response of focal individuals toward cues from social partners as well as on traits associated to cue emission in the social partners. We then included our surrogates of quality and quantity cue release in bivariate models of IGE, to test for correlations between DEE and IEE and DGE and IGE.

#### Direct genetic effects (DGE)

Univariate mixed effects models were used to quantify direct genetic effects (DGE) by partitioning the phenotypic variance (*V*_*P*_) of each behavioral trait (RA, RMV, ROT) and a composite trait made of the principal component axis that compiles all three traits (PC1) into: variation among sires (***V***_***S***_), among dams (***V***_***D***_) and within full-sib families (***V***_***W***_).

Following (Brommer 2013), and since we only had two environments (and thus two possible states per family) we took a character state approach and included feeding treatment as a numerical fixed effect variable (poor = 1 *vs* rich = 3), and as a categorical variable (poor *vs* rich) for random slopes. We also tested whether age, maximum carapace width, maximum abdomen width, weight loss and total recording time affected the variance partitioning of the traits under consideration, by running additional models with or without these variables as covariates. The overall model was:

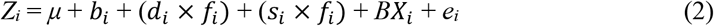

where Z_*i*_ is the behavioral trait of the focal individual *i, μ* is the mean of the trait (intercept), *b*_*i*_ the deviation from the block effect for individual *i, d*_*i*_ the deviation due to the dam family of individual *i, s*_*i*_ the deviation due to the sire family of the same individual, *f*_*i*_ the developmental feeding treatment of individual *i, X*_*i*_ is the vector of covariates of individual *i* and *B* the vector of covariate partial regression coefficients. Note that this matrix includes the coefficient for the feeding treatment as a fixed factor as well, which serves to accommodate the character state approach of phenotypic plasticity (Brommer 2013). Finally, *e*_*i*_ is the random deviation from causes other than the explanatory terms.

Narrow sense heritability (*h*^*2*^) was then estimated as the proportion of four times the sire variance component to the total phenotypic variance (*4V*_*S*_ */ V*_*P*_) and broad-sense heritability (*H*^*2*^) as the proportion of four times the dam variance to the total phenotypic variance (*4V*_*d*_ */ V*_*P*_). Common environmental effects were minimized by separating the offspring from their mothers relatively soon after they were born and including them in the split-brood feeding treatment (Henriques et al. 2021).

#### Indirect genetic and environmental effects (IGE and IEE)

We estimated the effect of social partners traits on traits in focal individuals with a combination of the variance-components (Griffing 1967; Bijma 2014) and the trait-based approach (Moore et al. 1997; McGlothlin and Brodie 2009), to estimate *r*_*DGE−IGE*_; i.e., DGE-IGE correlations (Costa E Silva et al. 2017; Santostefano et al. 2017*a*). These models allow partitioning the correlations between traits in focal individuals and in social partners into their genetic and environmental variance and covariance components. Bivariate models were built by pairing each behavioral trait recorded (RA, RMV, ROT, PC1) with each of its social partner traits hypothesized to mediate social interactions through conspecific cues (weight loss and body size). Using a variance partitioning approach, we decomposed the observed correlation between focal and social partner traits (i.e. the interaction coefficient) into genetic and environmental correlations (Garant et al. 2008; Wilson et al. 2010), as follows:

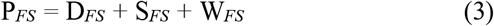

where P_*FS*_ is the phenotypic variance-covariance matrix of focal (*F* - behavioral) and social partner (*S* - morphological or physiological) individual traits, D_*FS*_ and S_*FS*_ the genetic (dam and sire) variance-covariance matrices of the same traits, and W_*FS*_ the within full-sib family variance-covariance matrix. Bivariate MCMCglmm models were then fitted with sire and dam of social partners (SIRE_SP_, DAM_SP_) as random effects and feeding treatment (FT_SP_), age (A_SP_) and carapace width (CW_SP_) of social partners as covariates. The latter being solely used in the bivariate models correlating weight loss. With this approach, we are also able to estimate correlations between direct (DEE) and indirect (IEE) environmental factors as the corresponding environmental (residual) components of the correlations estimated through variance partitioning (*r*_*DGE−IGE*_). Note that these *r*_*DGE−IGE*_ estimates differ from classical IGE estimates in that we include the dam component. This was done because the sire component of traits in social partners was not significant. We thus decided to distinguish only between non-additive genetic and maternal effects and purely environmental effects. To obtain the above variance-covariance components in a bivariate model, the two sub-models for the focal (*F*) and social partner (*S*) individuals were respectively:

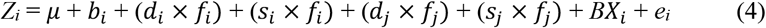

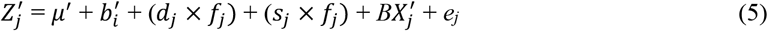

where *i* and *j* refer to the focal and social partner individuals respectively, and the ′s refer to traits and covariates of the social partner. All other terms are as in eq. 2.

#### Statistical analyses

All analyses were performed using R v.4.0.2 (R development Core Team 2020). To estimate variance and covariance components for each trait and the PC, we performed mixed effects animal models in the Bayesian framework of “MCMCglmm” (v.2.29) (Hadfield 2010). In all full models, dam and sire were included as random factors to account for the paternal half-sib/full-sib design structure. To make the paternal half-sib/full-sib design more explicit throughout the model (e.g., we did not measure traits over more than one single generation), we ran all final models including the identity of the sire as a random effect instead of using the pedigree option (ped) in MCMCglmm. All models were run twice to obtain two chains, and chain convergence was tested using the Gelman-Rubin diagnostic (Gelman and Rubin 1992), which was always met (i.e., close to 1 to the nearest 0.01 units). We set a burn in of 5000 samples and ran all models for sufficient iterations to ensure a minimum of 1950 effective samples for parameter estimation. Prior variances were set by dividing the phenotypic variance by the number of random effects (Wilson et al. 2010), while prior covariances were set to zero. A relatively low degree of belief (nu) was set at 0.2 for univariate models, at 2 for bivariate and random intercepts and slopes models and at 3 for three-variate models (genetic correlations). Sensitivity analyses using higher nu values (up to 1) for the univariate models revealed no substantial qualitative differences (not shown). Models with different fixed or random effects components were first compared using the Deviance Information Criterion (DIC), analogous to AIC (Burnham et al. 2011). A lower DIC value indicates a better fit between model and data (Li et al. 2017). As there is no standard DIC difference value between models for which one model should be chosen over another, models with the lowest DIC values, and with differences with the previous model of more than two units were tested for significance by comparing them with models including fewer parameters (e.g., only the block effect for trials performed in different clusters-times) or null models (with only the intercept), by means of mixture likelihood-ratio tests, using 0 and 1 degrees of freedom for univariate models, and 1 and 2 for bivariate models (Visscher 2006). Since the MCMCglmm algorithm calculates deviances and not log-likelihoods, to perform Likelihood ratio tests (LRTs) we re-implemented the models using the functions “glm” and “lmer”, the latter from the library “lme4” (Bates et al. 2007). We first compared a model including no random effects (null model) to another including only the block effect. If we found that the ΔDIC decreased and the mixture LRT was significant (e.g., >1.92 for univariate models with 0 and 1 d.f. and > 4.92 for bivariate models), then we kept the block effect and all other models (with dam and/or sire and/or the interaction with the feeding regime and/or the covariates) were compared to this one. Otherwise, all other models were compared to the null model. Posterior credible intervals (CI) for the estimates of narrow and broad-sense heritabilities, and genetic and maternal correlations were calculated from the posterior distributions using the highest-posterior-density function (HPD interval, package MCMCglmm (Hadfield 2010). Covariances were supported when 95% CI excluded zero. However, only variance components and heritabilities were reported for final models; i.e., when the model with sire and/or dam random effects had lower DIC values than null (or block) models, and the mixture LRT comparing them was significant. Reporting results of non-significant models (e.g., those including sire even though their DIC value is higher) may be misleading because estimates may be highly inflated depending on the degree of belief (the “nu” parameter in MCMCglmm). Because variances are bounded above zero, support of variances estimates was assessed by comparing the DIC values between fitted models.

## Results

### Direct Genetic Effects (DGEs)

The behavioural traits of the focal individual showed high variation among individuals, with means close to 0 and very wide ranges (fig. S1). However, individuals tended to avoid the side of the experimental arena that contained conspecific cues (relative occupancy time was -0.137 ± 0.533). This trait also showed significant broad sense heritability (*H*^*2*^ = 0.274 [CI: 0.071 to 0.526]), unlike the other behavioral traits. Also, no trait showed significant narrow-sense heritability, hence there were no DGE for these traits (Table 1). Additionally, we found significant phenotypic correlations (*r*_*p*_) among behavioural traits, namely a negative correlation between relative occupancy time and relative mean velocity (*r*_*p*_ = -0.548 [CI: -0.610; -0.486]), a negative correlation between relative occupancy time and relative activity (*r*_*p*_ = -0.432 [CI: -0.500; -0.358]) and a weaker positive correlation between relative mean velocity and relative activity (*r*_*p*_ = 0.232 [CI: 0.140; 0.300]). These correlations, however, were only environmental, not genetic, as revealed by comparing DIC values of multivariate MCMCglmm models with or without the dam and/or sire identity, which always presented higher DIC values than the model without these components (ΔDIC range 12-30). Moreover, by means of a principal component analysis (PCA), we found that ROT, RMV and RA were all grouped in one principal component axis explaining 61% of the total variance (fig. S2). From this PCA, we extracted the values of the first principal component (PC1), which represents the composite behaviour formed by these three traits. Given the loadings, individuals with a high PC1 score corresponded to animals that displayed higher activity and velocity in the patch with conspecific cues, but that visited that patch for shorter amounts of time. This composite behaviour also revealed no DGE (Table 1). Moreover, we did not find signs of Genotype x Environment (GxE) interactions, nor of the effect of the feeding treatment *per se*, on any of the behavioral traits (Table 1), indicating that food availability during development had little effect on their behaviour toward conspecific cues. Similarly, including the covariates age, total time recorded, body size, body condition and weight loss of the focal individuals did not improve model fits (Table 1).

**Table 1.**
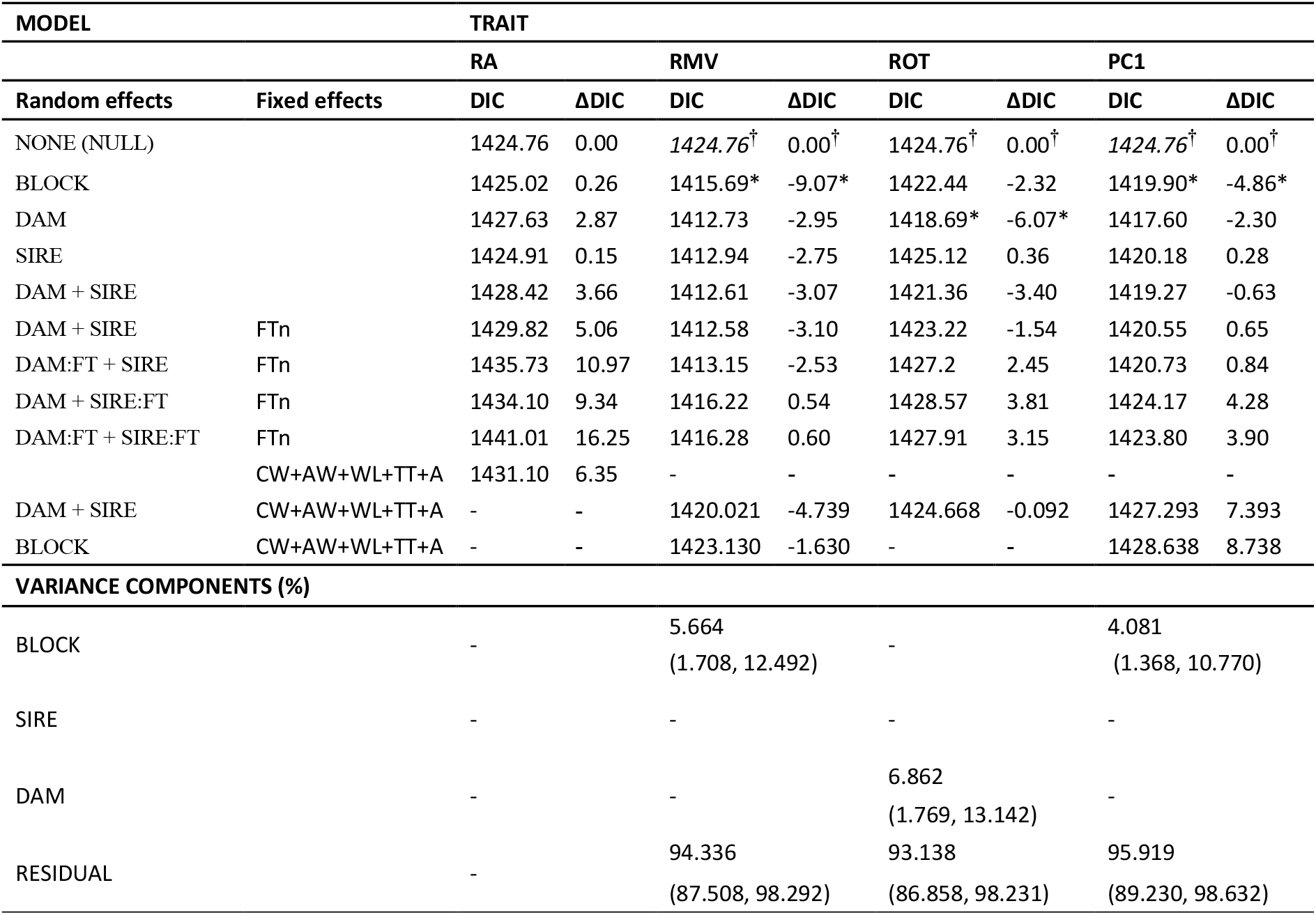
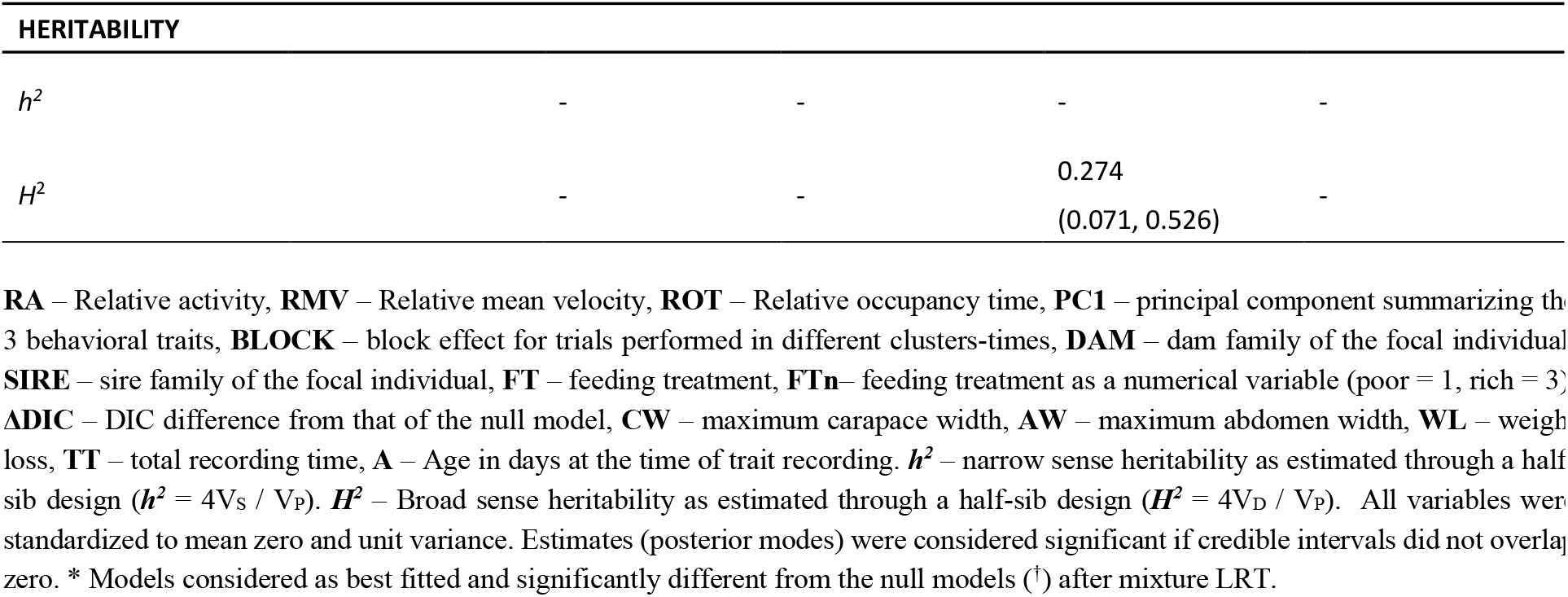
Direct genetic effects (DGEs) on behaviors as estimated from a half-sib analysis in univariate Bayesian GLMMs (N = 563).

| MODEL |  | TRAIT |  |  |  |  |  |  |  |
| --- | --- | --- | --- | --- | --- | --- | --- | --- | --- |
|  |  | RA |  | RMV |  | ROT |  | PC1 |  |
| Random effects | Fixed effects | DIC | ΔDIC | DIC | ΔDIC | DIC | ΔDIC | DIC | ΔDIC |
| NONE (NULL) |  | 1424.76 | 0.00 | 1424.76<sup>†</sup> | 0.00<sup>†</sup> | 1424.76<sup>†</sup> | 0.00<sup>†</sup> | 1424.76<sup>†</sup> | 0.00<sup>†</sup> |
| BLOCK |  | 1425.02 | 0.26 | 1415.69* | -9.07* | 1422.44 | -2.32 | 1419.90* | -4.86* |
| DAM |  | 1427.63 | 2.87 | 1412.73 | -2.95 | 1418.69* | -6.07* | 1417.60 | -2.30 |
| SIRE |  | 1424.91 | 0.15 | 1412.94 | -2.75 | 1425.12 | 0.36 | 1420.18 | 0.28 |
| DAM + SIRE |  | 1428.42 | 3.66 | 1412.61 | -3.07 | 1421.36 | -3.40 | 1419.27 | -0.63 |
| DAM + SIRE | FTn | 1429.82 | 5.06 | 1412.58 | -3.10 | 1423.22 | -1.54 | 1420.55 | 0.65 |
| DAM:FT + SIRE | FTn | 1435.73 | 10.97 | 1413.15 | -2.53 | 1427.2 | 2.45 | 1420.73 | 0.84 |
| DAM + SIRE:FT | FTn | 1434.10 | 9.34 | 1416.22 | 0.54 | 1428.57 | 3.81 | 1424.17 | 4.28 |
| DAM:FT + SIRE:FT | FTn | 1441.01 | 16.25 | 1416.28 | 0.60 | 1427.91 | 3.15 | 1423.80 | 3.90 |
|  | CW+AW+WL+TT+A | 1431.10 | 6.35 | - | - | - | - | - | - |
| DAM + SIRE | CW+AW+WL+TT+A | - | - | 1420.021 | -4.739 | 1424.668 | -0.092 | 1427.293 | 7.393 |
| BLOCK | CW+AW+WL+TT+A | - | - | 1423.130 | -1.630 | - | - | 1428.638 | 8.738 |
| **VARIANCE COMPONENTS (%)** |  |  |  |  |  |  |  |  |  |
| BLOCK |  | - |  | 5.664<br>(1.708, 12.492) |  | - |  | 4.081<br>(1.368, 10.770) |  |
| SIRE |  | - |  | - |  | - |  | - |  |
| DAM |  | - |  | - |  | 6.862<br>(1.769, 13.142) |  | - |  |
| RESIDUAL |  | - |  | 94.336<br>(87.508, 98.292) |  | 93.138<br>(86.858, 98.231) |  | 95.919<br>(89.230, 98.632) |  |

**HERITABILITY**
|  |  |  |  |  |
| --- | --- | --- | --- | --- |
| $h^2$ | - | - | - | - |
| $H^2$ | - | - | 0.274<br>(0.071, 0.526) | - |

**RA** – Relative activity, **RMV** – Relative mean velocity, **ROT** – Relative occupancy time, **PC1** – principal component summarizing the 3 behavioral traits, **BLOCK** – block effect for trials performed in different clusters-times, **DAM** – dam family of the focal individual, **SIRE** – sire family of the focal individual, **FT** – feeding treatment, **FTn**– feeding treatment as a numerical variable (poor = 1, rich = 3), **ΔDIC** – DIC difference from that of the null model, **CW** – maximum carapace width, **AW** – maximum abdomen width, **WL** – weight loss, **TT** – total recording time, **A** – Age in days at the time of trait recording. $h^2$ – narrow sense heritability as estimated through a half-sib design ( $h^2 = 4V_S / V_P$ ). $H^2$ – Broad sense heritability as estimated through a half-sib design ( $H^2 = 4V_D / V_P$ ). All variables were standardized to mean zero and unit variance. Estimates (posterior modes) were considered significant if credible intervals did not overlap zero. \* Models considered as best fitted and significantly different from the null models (†) after mixture LRT.

Concerning traits associated to cue release in the social partners, after controlling for the age of the individuals, carapace width showed high broad sense heritability (dam only model) and substantial evidence for GxE as estimated from the interaction between dam and feeding treatment (Table 2). Broad sense heritability estimates were substantially higher in rich environments. Weight loss also showed substantial broad sense heritability (although on the sire only model) after controlling for feeding treatment, carapace width and age, all of which affected the rate of weight loss. No sign of GxE was found (Table 2). As expected, older spiders and spiders in the rich feeding treatment were larger. Also, a rich environment made spiders losing less weight, while larger and older spiders lost more weight (Table 2).

**Table 2.**
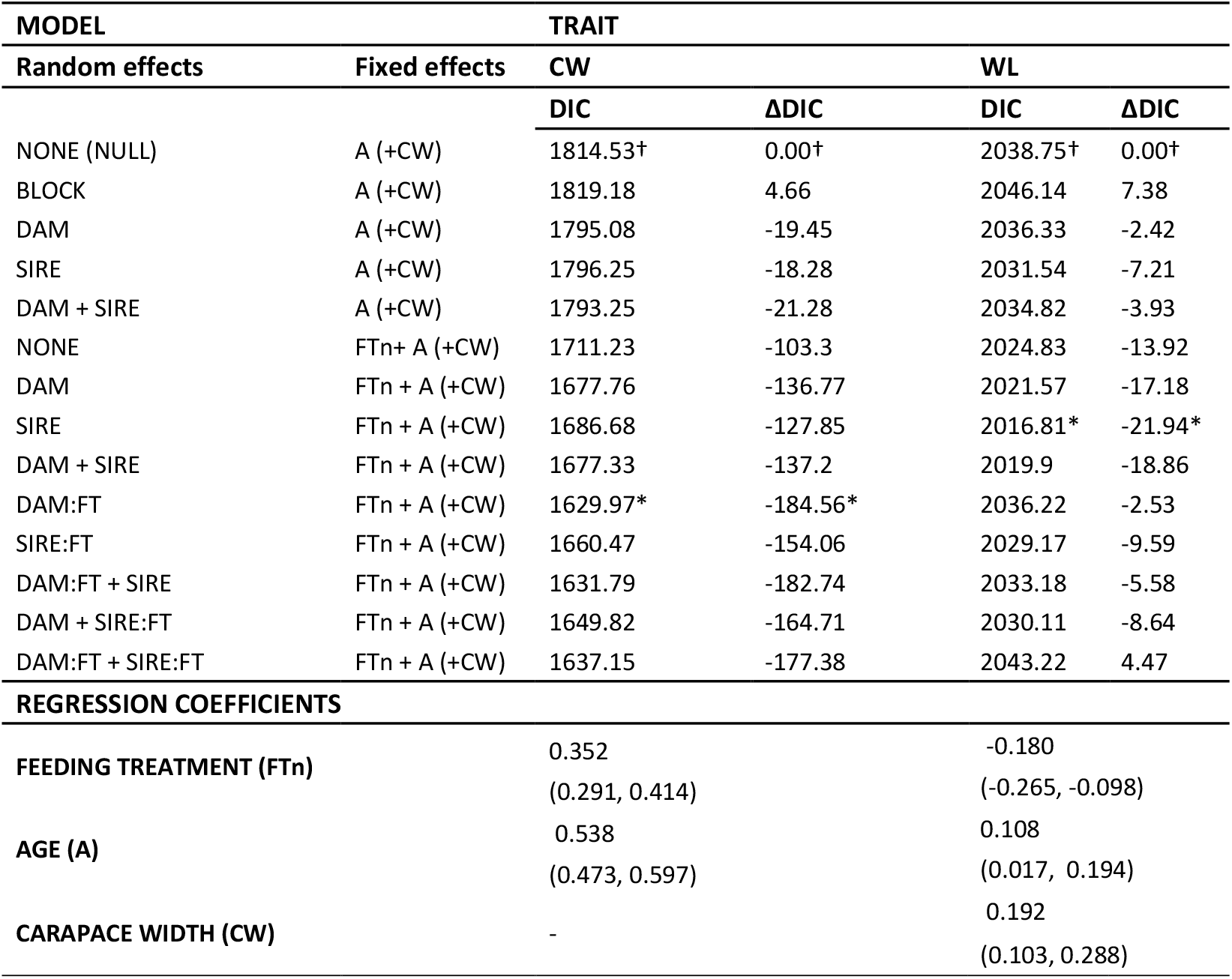
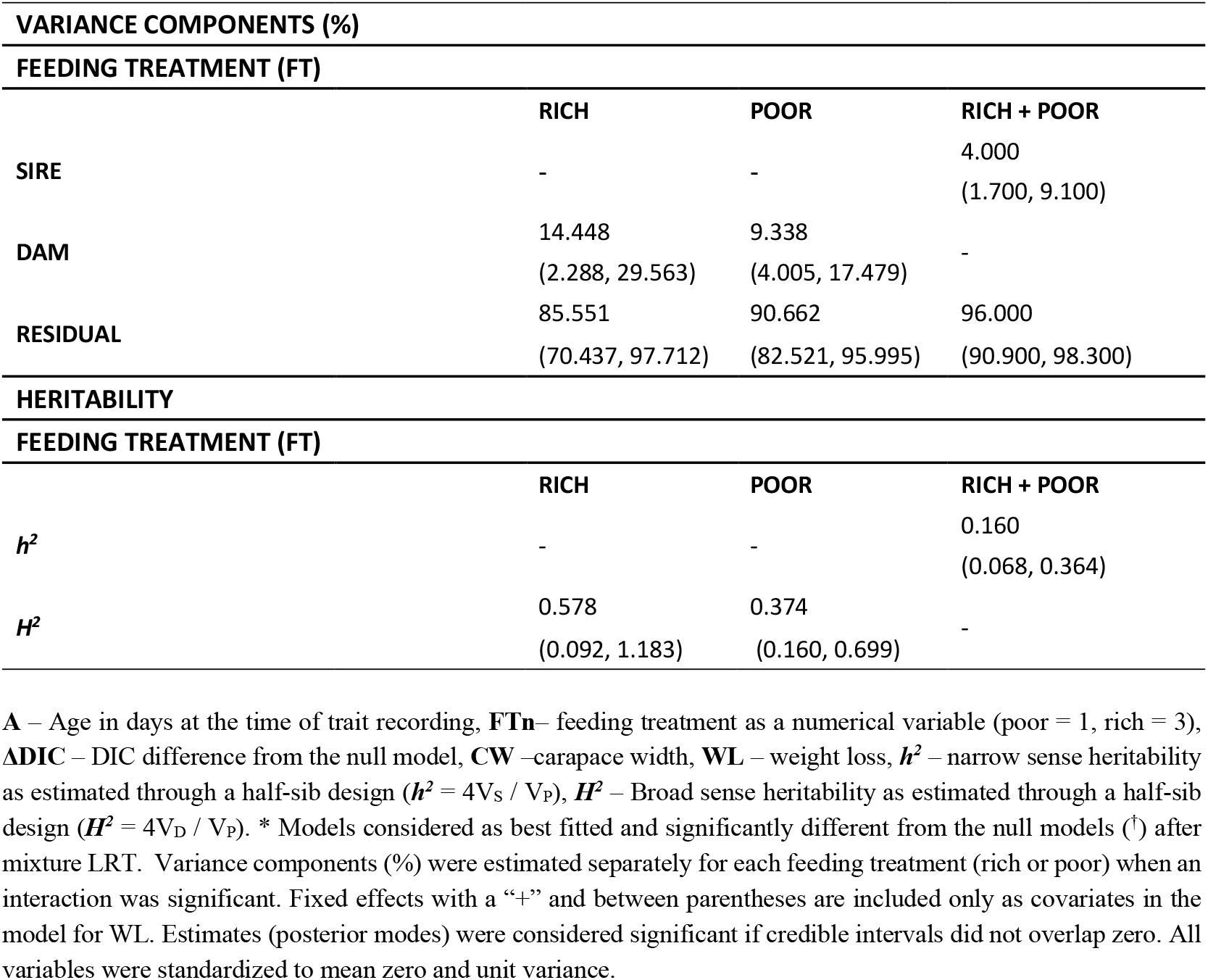
Direct genetic effects (DGEs) on carapace width (CW) and weight loss (WL) as estimated from a half-sib analysis and univariate Bayesian GLMMs (N = 728). Estimates were calculated for each feeding treatments separately when there was a significant evidence of genotype-by-environment (GxE) interactions.

### DGE-IGE correlations

For weight loss, the bivariate approach failed to find among-sire components for DGE-IGE correlations. Instead, the correlations were totally ascribed to the within full-sib component (DEE-IEE) for all behavioural traits. In absolute values, the coefficient ranged from 0.14 to 0.20 (Fig. 1). In general, according to the negative correlation for the composite PC1 (i.e., negative DEE-IEE correlation), social partners that lost more weight from environmental causes elicited responses on focal animals that involved spending relatively more time in patches with conspecific cues and moving relatively less frequently and more slowly in those patches.

**Figure 1.**
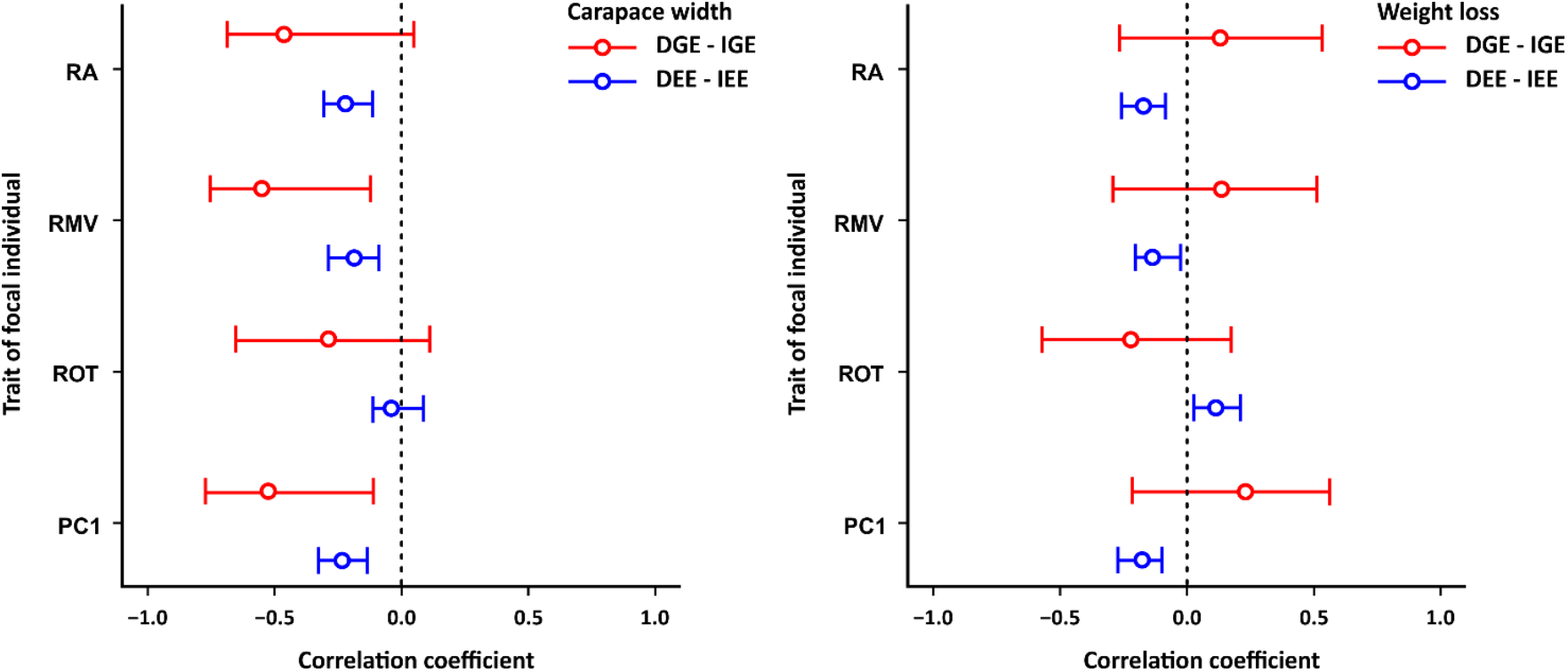
Correlation coefficients for maximum carapace width (left panel) and for weight loss (right panel) in the social partner and behavioral traits in focal individuals. Correlations between genetic and indirect genetic effects (DGE-IGE). Significant estimates do not overlap zero (dashed line). **RA** – relative activity; **RMV** – relative mean velocity; **ROT** – relative occupancy time; **PC1** – composite behavior.

The bivariate approach detected significant and moderately strong negative DGE-IGE correlation coefficients among dams (range of the absolute *r* value for DGE-IGE: 0.52-0.55, fig. 1; Table 3) for carapace width on relative mean velocity and PC1, being the within full-sib (environmental – DEE-IEE) coefficients substantially lower (range of the absolute *r* value: 0.18 -0.23, fig.1; Table 3). For relative activity, we found only negative environmental correlated effects for the social partners’ body size (Fig. 1, Table 3). Thus, social partners that were larger (due to either common environmental, dominance or maternal effects contained in the dam component) elicited focal individuals to spend more time in patches with conspecific cues but to move less and more slowly in those patches.

**Table 3.** DGE-IGE and DEE-IEE correlation models between morphological and physiological traits and behaviors as estimated from (co)variances in a half-sib analysis fitting multivariate Bayesian GLMMs (N = 563).

| MODEL |  | FOCAL TRAIT |  |  |  |  |  |  |  |  |
| --- | --- | --- | --- | --- | --- | --- | --- | --- | --- | --- |
| Random effects | Fixed effects | Trait | RA |  | RMV |  | ROT |  | PC1 |  |
|  |  |  | DIC | ΔDIC | DIC | ΔDIC | DIC | ΔDIC | DIC | ΔDIC |
|  |  | CW<sub>SP</sub> |  |  |  |  |  |  |  |  |
| NONE (NULL) | A<sub>SP</sub> |  | 2740.71<sup>†</sup> | 0.00<sup>†</sup> | 2843.87<sup>†</sup> | 0.00<sup>†</sup> | 2696.91<sup>†</sup> | 0.00<sup>†</sup> | 2725.99<sup>†</sup> | 0.00<sup>†</sup> |
| DAM<sub>SP</sub> | A<sub>SP</sub> |  | 2709.38 | -31.321 | 2712.43 | -131.46 | 2695.92 | -1.00 | 2697.43 | -28.56 |
| DAM<sub>SP</sub> | FTn<sub>SP</sub> + A<sub>SP</sub> |  | 2709.38 | -31.321 | 2687.15 | -156.74 | 2662.52 | -34.39 | 2663.99* | -62.00* |
| DAM<sub>SP</sub>:FT<sub>SP</sub> | FTn<sub>SP</sub> + A<sub>SP</sub> |  | 2695.89* | -44.814* | 2645.54* | -198.34* | 2656.34* | -40.57* | 2666.42 | -59.56 |
| DAM<sub>SP</sub>:FT<sub>SP</sub> + SIRE<sub>SP</sub>:FT<sub>SP</sub> | FTn<sub>SP</sub> + A<sub>SP</sub> |  | 2704.85 | -35.855 | 2737.06 | -106.82 | 2665.35 | -31.56 | 2675.02 | -50.97 |
|  |  | WL<sub>SP</sub> |  |  |  |  |  |  |  |  |
| NONE (NULL) |  |  | 2837.08<sup>†</sup> | 0.00<sup>†</sup> | 2843.36<sup>†</sup> | 0.00<sup>†</sup> | 2840.25<sup>†</sup> | 0.00<sup>†</sup> | 2837.25<sup>†</sup> | 0.00<sup>†</sup> |
| NONE | FTn<sub>SP</sub> + CW<sub>SP</sub> + A<sub>SP</sub> |  | 2832.63* | -4.455* | 2829.50* | -13.85* | 2813.36* | -26.89* | 2832.65 | -4.59 |
| SIRE<sub>SP</sub> |  |  | 2847.80 | 10.719 | 2856.32 | 12.96 | 2843.73 | 3.48 | 2840.00 | 2.76 |
| SIRE<sub>SP</sub> | FTn<sub>SP</sub> + CW<sub>SP</sub> + A<sub>SP</sub> |  | 2841.51 | 4.425 | 2836.69 | -6.66 | 2817.43 | -22.82 | 2831.70* | -5.55* |

**RA** – Relative activity, **RMV** – Relative mean velocity, **ROT** – Relative occupancy time, **PC1** – principal component holding the 3 behavioral traits, **FT** – feeding treatment, **DAM** – dam family of the focal individual, **SIRE** – sire family of the focal individual, **DAM<sub>SP</sub>** – dam family of social partner, **SIRE<sub>SP</sub>** – sire family of social partner, **FT<sub>SP</sub>** –feeding treatment of social partner, **FTn<sub>SP</sub>** –feeding treatment of social partner as a numerical variable (low=1, high=3), **A<sub>SP</sub>** - Age of social partner, **CW<sub>SP</sub>** – maximum carapace width of social partner, **AW<sub>SP</sub>** – abdomen width of social partner, **WL<sub>SP</sub>** – weight loss of social partner, **ΔDIC** – DIC difference from the lowest DIC model. All variables were standardized to unit variance. \* Models considered as best fitted and significantly different from the null, or second best models (†) after mixture LRT. Interaction coefficients ( $\psi$ ) were considered significant if credible intervals do not overlap 0.

## Discussion

In this study, we show that spiders tended to avoid patches previously occupied by conspecifics. However, this social response was highly variable, with some individuals spending a high proportion of time in patches with conspecific cues. Additionally, individuals that spent more time in those patches moved relatively less often and at relatively lower speed. However, we found no significant additive genetic variance for any trait, and only a significant broad-sense heritability for relative occupancy time. We also found evidence for such broad-sense heritability in traits associated with the production of cues, body size and weight loss. These traits were also genetically (non-additive IGE) and/or environmentally (IEE) correlated with behavioural traits in focal individuals. Below we discuss the ecological and evolutionary implications of these findings.

Focal individuals tended to avoid patches with conspecific cues on average, but they spent more time in those patches when the social partners were larger or had lost more weight. This is intriguing because this behaviour increases their risk of being cannibalized and the latter increases with differences in body size (Hvam et al. 2005; Rypstra and Samu 2006; Wise 2006; Gonzalez 2012). Thus, assuming that the size of focal individuals is randomly distributed across trials, this behaviour entails that they are often entering patches where their risk of being cannibalized is high. This behaviour calls for an explanation. One possibility is that focal individuals are attracted to patches where the benefits of feeding on a conspecific are higher. Alternatively, spiders could associate conspecific cues to patches where the chances to find prey are high (Valone and Templeton 2002). Finally, it is possible that, by not including social partners themselves and their behaviour in our experimental design, we limited the amount of cues accessible to focal individuals.

Although the average value of most traits did not differ from null expectations, their variance was not randomly distributed across trait space (fig. 2). Indeed, we found that trait values were distributed along a principal component axis, where individuals that spent more time on patches with conspecific cues (higher relative occupancy) moved less and slower on those patches (lower relative activity and velocity). This is at odds with the usual relationship among these behavioural traits, in which individuals that move at a faster rate also tend to explore novel environments more often. Possibly, this difference pertains to the fact that the ‘classical’ boldness-exploratory syndrome is generally identified by placing individuals in patches with either predators or prey. Bold individuals tend to be more effective foragers (Maskrey et al. 2018), but at the expenses of being more often detected by their own predators. In our case, because we are dealing with a cannibalistic interaction, the same individual may be a predator or a prey. We therefore identified a spectrum of social interactions with two different strategies at the extremes: careful sneakers and careless avoiders (fig. 2c). Both these strategies are likely to result in individuals being less attacked by conspecifics. Indeed, avoiding patches with conspecific cues is an efficient way to avoid agonistic interactions. Additionally, because wolf spiders are very efficient at detecting movement (Rovner 1996), moving slower is likely to reduce conspicuousness towards conspecifics. Lower conspicuousness in the presence of conspecific cues can also be a hunting strategy to avoid being detected by potential conspecific prey. The actual existence of these strategies could be documented with further research testing the fitness consequences of the spectrum of behaviors along the PCA axis.

**Figure 2.**
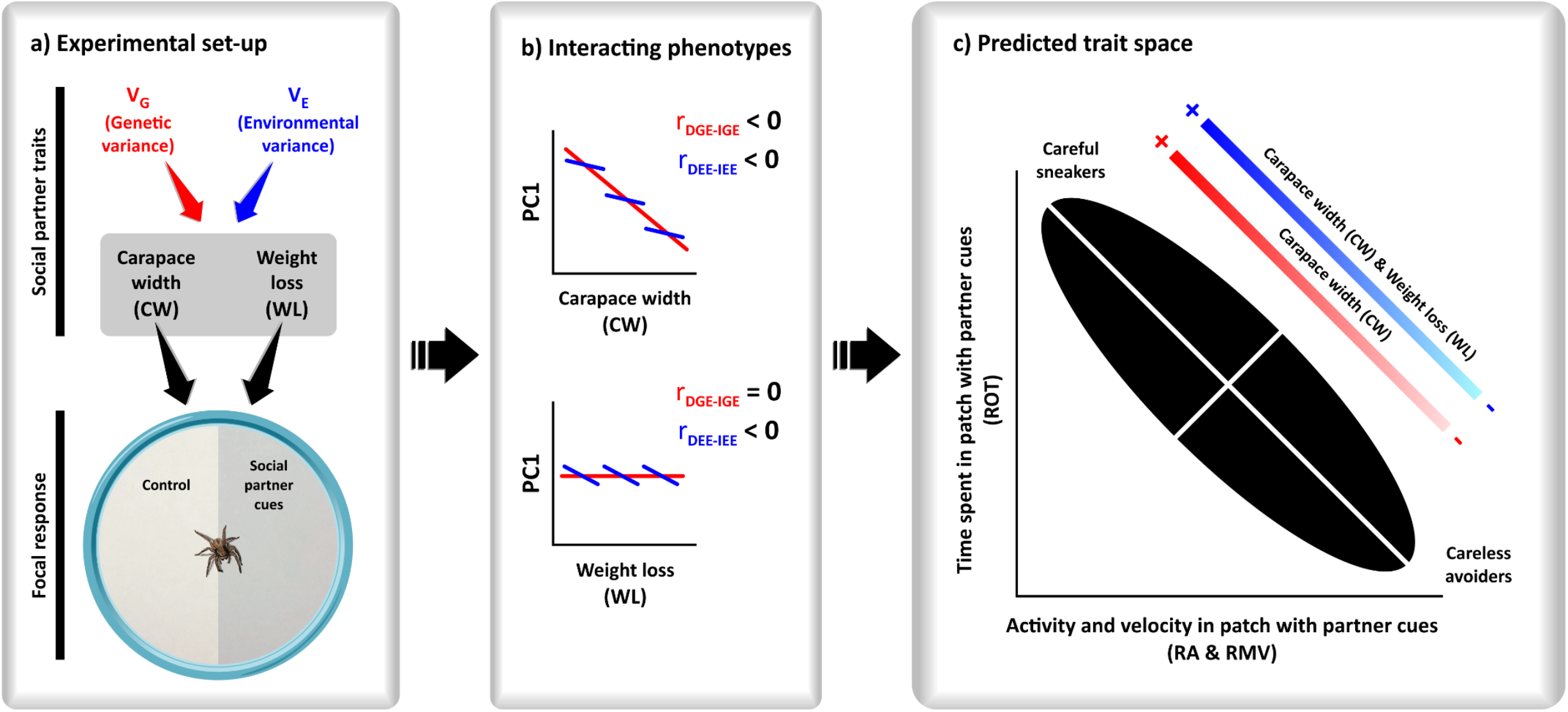
Schematic representation of the experimental design and main conclusions. **a)** Experimental arenas, in which spiders were free to travel between patches without (control) or with social partner cues. Weight loss, a trait measured in social partners, is predicted to be positively related to the amount of cues released. Body size is predicted to be related to the quality of the cues released; **b)** PC1 is a composite behaviour of focal individual relative activity (RA), mean velocity (RMV) and occupancy time (ROT) in/of patches with conspecific cues. A higher value of PC1 means animals moving at relatively higher speeds and relatively more often in the patch with conspecifics, but expending relatively less time in those patches (**fig. S2**). Both weight loss and carapace width show environmental correlations with PC1, and carapace width a genetic correlation. The different blue lines refer to within full-sib family (environmental) correlations. Only three maternal full-sib families have been depicted for simplicity; **c)** the magnitude of the genetic (red) or environmental (blue) correlations of weight loss and body size of social partners with the PC1 response of the focal individuals determine the strategy of the focal individual. For instance, lower genetic and environmental CW values in the social partner, induces focal individuals to follow a strategy closer to moving fast in conspecific patches but visiting them less often (careless avoider). Higher social partner CW values on the other hand, induce focal individuals to visit these patches more often but moving less frequently and at lower speeds once there (careful sneakers).

We failed to find any evidence for DGE on the principal component lumping all behavioural traits. However, we did find evidence for body size and weight loss in social partners affecting most behavioural traits in focal individuals, via IEE and non-additive IGE. Indeed, we found negative non-additive DGE-IGE correlations between carapace width and the principal component, with similar consistent relationships for the behavioral traits that underlied this composite trait. Moreover, we documented a DEE-IEE correlation between carapace width and weight loss with some of the behavioural traits in focal individuals, including the principal component axis. Because we found no evidence for additive genetic variance in traits in the social partners, we can exclude the potential for IGE to affect trait evolution in focal individuals. Still, traits in social partners affected behavioural traits in different directions, as well as the composite trait lumping all of these traits. Therefore, variation in traits in social partners (body size and weight loss), either due to broad-sense heritability or environmetal effects, may contribute to the maintenance of a continuum of strategies to deal with cannibalistic conspecifics in populations. These correlations also mean that not all strategies are available in the phenotypic space.

In conclusion, we unraveled a composite/complex behavioral phenotype that whose variability was determined by the environmental and non-additive genetic components of interacting conspecific phenotypes. We show that these behavioral traits may be correlated in ways that differ from those generally identified in other interactions, and that such correlation is determined by the social environment. Therefore, even in the absence of genetic constrains, organisms may not have access to the entire phenotypic space of behavioral traits. Non-additive IGE and IEE appear as important sources contributing to maintain a diversity of behavioral strategies in the wild.

## Acknowledgements

We thank D. Mayntz and K. Jensen for providing the recipes for the media used to culturing nitrogen-enriched Drosophila. This study was supported by a PhD grant (PD/BD/106059/2015) attributed to JFH by the Portuguese Science and Technology foundation (FCT), by the grant P12-RMN-1521 from the Andalusian government and by the grant CGL2015-66192-R from the Spanish Ministry of Economy and Competitiveness, both partially funded by the European Regional Development Found, both attributed to Jordi Moya-Laraño and to the FPU scholarship (FPU13/04933) from the Spanish Ministerio de Educacion, Cultura y Deporte to Dolores Ruiz-Lupión.

## References

Arnqvist, G., and S. Henriksson. 1997. Sexual cannibalism in the fishing spider and a model for the evolution of sexual cannibalism based on genetic constraints. Evolutionary Ecology 11:255–273.

Bailey, N. W., L. Marie-Orleach, and A. J. Moore. 2018. Indirect genetic effects in behavioral ecology: does behavior play a special role in evolution? Behavioral Ecology 29:1–11.

Bailey, N. W., and M. Zuk. 2012. Socially flexible female choice differs among populations of the pacific field cricket: Geographical variation in the interaction coefficient psi (ψ). Proceedings of the Royal Society B: Biological Sciences 279:3589– 3596.

Barnes, M. C., M. H. Persons, and A. L. Rypstra. 2002. The effect of predator chemical cue age on antipredator behavior in the wolf spider Pardosa milvina (Araneae: Lycosidae). Journal of Insect Behavior 15:269–281.

Barrientos, J. A. 2004. Lycosa Ambigua sp. Nov.(Araneae, Lycosidae), una nueva tarántula para la fauna ibérica. Revista ibérica de aracnología 9:23–29.

Bates, D., D. Sarkar, M. Bates, and L. Matrix. 2007. The lme4 Package (R Package, Version 2).

Bijma, P. 2014. The quantitative genetics of indirect genetic effects: A selective review of modelling issues. Heredity 112:61–69.

Blake, C. A., M. L. Andersson, K. Hulthén, P. A. Nilsson, and C. Brönmark. 2018. Conspecific boldness and predator species determine predation-risk consequences of prey personality. Behavioral Ecology and Sociobiology 72:1–7.

Bleakley, B. H., and E. D. Brodie Iv. 2009. Indirect genetic effects influence antipredator behavior in guppies: Estimates of the coefficient of interaction psi and the inheritance of reciprocity. Evolution 63:1796–1806.

Bleakley, B. H., S. M. Welter, K. McCauley-Cole, S. M. Shuster, and A. J. Moore. 2013. Cannibalism as an interacting phenotype: precannibalistic aggression is influenced by social partners in the endangered Socorro Isopod (Thermosphaeroma thermophilum). Journal of evolutionary biology 26:832–842.

Bonnet, T., M. B. Morrissey, P. de Villemereuil, S. C. Alberts, P. Arcese, L. D. Bailey, S. Boutin, et al. 2022. Genetic variance in fitness indicates rapid contemporary adaptive evolution in wild animals. Science 376:1012–1016.

Bonte, D., J. V. Borre, L. Lens, and J.-P. Maelfait. 2006. Geographical variation in wolf spider dispersal behaviour is related to landscape structure. Animal behaviour 72:655– 662.

Bonte, D., and L. Lens. 2007. Heritability of spider ballooning motivation under different wind velocities. Evolutionary Ecology Research 9:817–827.

Bonte, D., S. Van Belle, and J.-P. Maelfait. 2007. Maternal care and reproductive state-dependent mobility determine natal dispersal in a wolf spider. Animal Behaviour 74:63– 69.

Brommer, J. E. 2013. Variation in plasticity of personality traits implies that the ranking of personality measures changes between environmental contexts: Calculating the cross-environmental correlation. Behavioral Ecology and Sociobiology 67:1709–1718.

Brommer, J. E., and K. Rattiste. 2008. “Hidden” reproductive conflict between mates in a wild bird population. Evolution 62:2326–2333.

Burnham, K. P., D. R. Anderson, and K. P. Huyvaert. 2011. AIC model selection and multimodel inference in behavioral ecology: some background, observations, and comparisons. Behavioral ecology and sociobiology 65:23–35.

Carter, M. J., A. J. Wilson, A. J. Moore, and N. J. Royle. 2019. The role of indirect genetic effects in the evolution of interacting reproductive behaviors in the burying beetle, Nicrophorus vespilloides. Ecology and Evolution 9:998–1009.

Chakrabarty, A., P. Kronenberg, N. Toliopoulos, and H. Schielzeth. 2019. Direct and indirect genetic effects on reproductive investment in a grasshopper. Journal of Evolutionary Biology 32:331–342.

Costa e Silva, J., B. M. Potts, P. Bijma, R. J. Kerr, and D. J. Pilbeam. 2013. Genetic control of interactions among individuals: contrasting outcomes of indirect genetic effects arising from neighbour disease infection and competition in a forest tree. The New phytologist 197:631–641.

Costa E Silva, J., B. M. Potts, A. R. Gilmour, and R. J. Kerr. 2017. Genetic-based interactions among tree neighbors: Identification of the most influential neighbors, and estimation of correlations among direct and indirect genetic effects for leaf disease and growth in Eucalyptus globulus. Heredity 119:125–135.

Danielson-François, A. M., Y. Zhou, and M. D. Greenfield. 2009. Indirect genetic effects and the lek paradox: inter-genotypic competition may strengthen genotype × environment interactions and conserve genetic variance. Genetica 136:27–36.

Day, T., and R. Bonduriansky. 2011. A Unified Approach to the Evolutionary Consequences of Genetic and Nongenetic Inheritance. The American Naturalist 178:E18–E36.

Dewan, I., T. Garland, L. Hiramatsu, and V. Careau. 2019. I Smell a Mouse: Indirect Genetic Effects on Voluntary Wheel-Running Distance, Duration and Speed. Behavior Genetics 49:49–59.

Falconer, D., and T. Mackay. 1996. Introduction to quantitative genetics. Essex. UK: Longman Group.

Fernández-Montraveta, C., and J. Moya-Laraño. 2007. Sex-specific plasticity of growth and maturation size in a spider: implications for sexual size dimorphism. Journal of Evolutionary Biology 20:1689–1699.

Garant, D., J. D. Hadfield, L. E. Kruuk, and B. C. Sheldon. 2008. Stability of genetic variance and covariance for reproductive characters in the face of climate change in a wild bird population. Molecular Ecology 17:179–188.

Gavín-Centol, M. P., S. Kralj-Fišer, E. De Mas, D. Ruiz-Lupión, and J. Moya-Laraño. 2017. Feeding regime, adult age and sexual size dimorphism as determinants of pre-copulatory sexual cannibalism in virgin wolf spiders. Behavioral Ecology and Sociobiology 71.

Gelman, A., and D. B. Rubin. 1992. Inference from iterative simulation using multiple sequences. Statistical science 457–472.

Genung, M. A., J. K. Bailey, and J. A. Schweitzer. 2012. Welcome to the neighbourhood: interspecific genotype by genotype interactions in Solidago influence above- and belowground biomass and associated communities: Interspecific genotype × genotype interactions. Ecology Letters 15:65–73.

Germain, R. R., M. E. Wolak, P. Arcese, S. Losdat, and J. M. Reid. 2016. Direct and indirect genetic and fine-scale location effects on breeding date in song sparrows. Journal of Animal Ecology 85:1613–1624.

Gonzalez, D. N. 2012. The influence of size on cannibalism and predation in hungry wolf spiders (Lycosidae, Hogna crispipes).

Griffing, B. 1967. Selection in Reference to Biological Groups. Australian Journal of Biological Sciences 22:131.

Hadfield, J. D. 2010. MCMC methods for multi-response generalized linear mixed models: the MCMCglmm R package. Journal of statistical software 33:1–22.

Han, C. S., C. Tuni, J. Ulcik, and N. J. Dingemanse. 2018. Increased developmental density decreases the magnitude of indirect genetic effects expressed during agonistic interactions in an insect. Evolution 72:2435–2448.

Hanna, C., and P. Eason. 2013. Juvenile crab spiders (Mecaphesa asperata) use indirect cues to choose foraging sites. Ethology Ecology & Evolution 25:161–173.

Hendrickx, F., J. Maelfait, and L. Lens. 2008. Effect of metal stress on life history divergence and quantitative genetic architecture in a wolf spider. Journal of evolutionary biology 21:183–193.

Henriques, J. 2020. Intraspecific variation in functional traits in the wolf spider Lycosa fasciiventris: Implications for trophic cascades. PhD thesis, Biodiversity, Genetics and Evolution, University of Lisbon, Faculty of sciences, Lisbon, Portugal. Retrieved from the https://repositorio.ul.pt/handle/10451/44162

Henriques, J. F., M. Lacava, C. Guzmán, M. P. Gavín-Centol, D. Ruiz-Lupión, E. De Mas, S. Magalhães, et al. 2021. The sources of variation for individual prey-to-predator size ratios. Heredity 126:684–694.

Hunt, J., and L. W. Simmons. 2002. The genetics of maternal care: Direct and indirect genetic effects on phenotype in the dung beetle Onthophagus taurus. Proceedings of the National Academy of Sciences 99:6828–6832.

Hvam, A., D. Mayntz, and R. K. Nielsen. 2005. Factors Affecting Cannibalism Among Newly Hatched Wolf Spiders (Lycosidae, Pardosa Amentata). Journal of Arachnology 33:377–383.

Jensen, K., D. Mayntz, S. Toft, D. Raubenheimer, and S. J. Simpson. 2011. Nutrient regulation in a predator, the wolf spider Pardosa prativaga. Animal Behaviour 81:993– 999.

Kölliker, M., E. D. Brodie Iii, and A. J. Moore. 2005. The Coadaptation of Parental Supply and Offspring Demand. The American Naturalist 166:506–516.

Lane, S. M., A. J. Wilson, and M. Briffa. 2020. Analysis of direct and indirect genetic effects in fighting sea anemones. Behavioral Ecology 31:540–547.

Li, Y., Y. Jun, and T. Zeng. 2017. Deviance information criterion for Bayesian model selection: Justification and variation.

Lynch, M., and B. Walsh. 1998. Genetics and analysis of quantitative traits. (Vol. 1). Sinauer Sunderland, MA.

Marie-Orleach, L., N. Vogt-Burri, P. Mouginot, A. Schlatter, D. B. Vizoso, N. W. Bailey, and L. Schärer. 2017. Indirect genetic effects and sexual conflicts: Partner genotype influences multiple morphological and behavioral reproductive traits in a flatworm. Evolution 71:1232–1245.

Maskrey, D. K., S. J. White, A. J. Wilson, and T. M. Houslay. 2018. Who dares does not always win: risk-averse rockpool prawns are better at controlling a limited food resource. Animal Behaviour 140:187–197.

McGlothlin, J. W., and E. D. Brodie. 2009. How to measure indirect genetic effects: The congruence of trait-based and variance-partitioning approaches. Evolution 63:1785– 1795.

McGlothlin, J. W., A. J. Moore, J. B. Wolf, and E. D. Brodie Iii. 2010. Interacting phenotypes and the evolutionary process. III. Social evolution. Evolution 64:2558–2574.

Milanovich, J. R., and M. E. Hopton. 2016. Stoichiometry of Excreta and Excretion Rates of a Stream-dwelling Plethodontid Salamander. Copeia 2016:26–34.

Miller, C. W., and A. J. Moore. 2007. A potential resolution to the lek paradox through indirect genetic effects. Proceedings of the Royal Society B: Biological Sciences 274:1279–1286.

Mogali, S. M., S. K. Saidapur, and B. A. Shanbhag. 2020. Behavioral responses of tadpoles of Duttaphrynus melanostictus (Anura: Bufonidae) to cues of starved and fed dragonfly larvae. Phyllomedusa 19:93–98.

Moiron, M., Y. G. Araya-Ajoy, C. Teplitsky, S. Bouwhuis, and A. Charmantier. 2020. Understanding the Social Dynamics of Breeding Phenology: Indirect Genetic Effects and Assortative Mating in a Long-Distance Migrant. The American Naturalist 196:566–576.

Moore, A. J., E. D. Brodie Iii, and J. B. Wolf. 1997. Indirect Genetic Effects of Social Interactions. Evolution 51:1352–1362.

Moskalik, B., and G. W. Uetz. 2011. Female hunger state affects mate choice of a sexually selected trait in a wolf spider. Animal Behaviour 81:715–722.

Moya-Larano, J. 2002. Senescence and food limitation in a slowly ageing spider. Functional Ecology 734–741.

Moya-Laraño, J. 2011. Genetic variation, predator–prey interactions and food web structure. Philosophical Transactions of the Royal Society B: Biological Sciences 366:1425–1437.

Nentwig, W. 1986. Non-webbuilding spiders: prey specialists or generalists? Oecologia 69:571–576.

Persons, M. H., and A. L. Rypstra. 2000. Preference for chemical cues associated with recent prey in the wolf spider Hogna helluo (Araneae: Lycosidae). Ethology 106:27–35.

Persons, M. H., and A. L. Rypstra. 2001. Wolf spiders show graded antipredator behavior in the presence of chemical cues from different sized predators. Journal of Chemical Ecology 27:2493–2504.

Persons, M. H., S. E. Walker, A. L. Rypstra, and S. D. Marshall. 2001. Wolf spider predator avoidance tactics and survival in the presence of diet-associated predator cues (Araneae: Lycosidae). Animal Behaviour 61:43–51.

Pischedda, A., and A. K. Chippindale. 2006. Intralocus Sexual Conflict Diminishes the Benefits of Sexual Selection. PLoS Biology 4:e356.

Planas, E., C. Fernández-Montraveta, and C. Ribera. 2013. Molecular systematics of the wolf spider genus Lycosa (Araneae: Lycosidae) in the Western Mediterranean Basin. Molecular Phylogenetics and Evolution 67:414–428.

Rádai, Z., B. Kiss, and Z. Barta. 2017. Pace of life and behaviour: rapid development is linked with increased activity and voracity in the wolf spider Pardosa agrestis. Animal Behaviour 126:145–151.

Rebar, D., and R. L. Rodríguez. 2013. Genetic variation in social influence on mate preferences. Proceedings of the Royal Society B: Biological Sciences 280:20130803.

Ribeiro, D., A. R. Nunes, M. Teles, S. Anbalagan, J. Blechman, G. Levkowitz, and R. F. Oliveira. 2020. Genetic variation in the social environment affects behavioral phenotypes of oxytocin receptor mutants in zebrafish. Elife 9:e56973.

Roberts, J. A., and G. W. Uetz. 2004. Chemical signaling in a wolf spider: a test of ethospecies discrimination. Journal of Chemical Ecology 30:1271–1284.

Rode, N. O., P. Soroye, R. Kassen, and H. D. Rundle. 2017. Air-borne genotype by genotype indirect genetic effects are substantial in the filamentous fungus Aspergillus nidulans. Heredity 119:1–7.

Rovner, J. S. 1996. Conspecific interactions in the lycosid spider Rabidosa rabida: the roles of different senses. Journal of Arachnology 16–23.

Rudin, F. S., J. L. Tomkins, and L. W. Simmons. 2018. The effects of the social environment and physical disturbance on personality traits. Animal Behaviour 138:109– 121.

Rypstra, A. L., and F. Samu. 2005. Size Dependent Intraguild Predation and Cannibalism in Coexisting Wolf Spiders (Araneae, Lycosidae). Journal of Arachnology 33:390–397.

Rypstra, A. L., and F. Samu. 2006. Size Dependent Intraguild Predation and Cannibalism in Coexisting Wolf Spiders (Araneae, Lycosidae). Journal of Arachnology 33:390–397.

Santostefano, F., H. Allegue, D. Garant, P. Bergeron, and D. Réale. 2021. Indirect genetic and environmental effects on behaviors, morphology, and life-history traits in a wild Eastern chipmunk population. Evolution 75:1492–1512.

Santostefano, F., A. J. Wilson, P. T. Niemelä, and N. J. Dingemanse. 2017a. Indirect genetic effects: a key component of the genetic architecture of behaviour. Scientific Reports 7:10235.

Santostefano, F., A. J. Wilson, P. T. Niemelä, and N. J. Dingemanse. 2017b. Behavioural mediators of genetic life-history trade-offs: A test of the pace-of-life syndrome hypothesis in field crickets. Proceedings of the Royal Society B: Biological Sciences 284:15–17.

Schmitz, O. J., and K. B. Suttle. 2001. Effects of top predator species on direct and indirect interactions in a food web. Ecology 82:2072–2081.

Schrader, M., B. J. M. Jarrett, and R. M. Kilner. 2018. Parental care and sibling competition independently increase phenotypic variation among burying beetle siblings. Evolution 72:2546–2552.

Shannon, H., D. Kutz, and M. Persons. 2022. The effects of prenatal predator cue exposure on offspring substrate preferences in the wolf spider Tigrosa helluo. Animal Behaviour 183:41–50.

Shuster, S., E. Lonsdorf, G. Wimp, J. Bailey, and T. Whitham. 2006. Community heritability measures the evolutionary consequences of indirect genetic effects on community structure. Evolution 60:991–1003.

Storm, J. J., and S. L. Lima. 2010. Mothers forewarn offspring about predators: a transgenerational maternal effect on behavior. The American Naturalist 175:382–390.

Team, R. C. 2020. R: A language and environment for statistical computing (Version 4.0. 2). R Foundation for Statistical Computing.

Valone, T. J., and J. J. Templeton. 2002. Public information for the assessment of quality: A widespread social phenomenon. Philosophical Transactions of the Royal Society B: Biological Sciences 357:1549–1557.

Visscher, P. M. 2006. A note on the asymptotic distribution of likelihood ratio tests to test variance components. Twin research and human genetics 9:490–495.

Wetter, M. B., B. Wernisch, and S. Toft. 2012. Tests for attraction to prey and predator avoidance by chemical cues in spiders of the beech forest floor. Arachnologische Mitteilungen 84–89.

Whitlock, R., M. C. Bilton, J. P. Grime, and T. Burke. 2011. Fine-scale community and genetic structure are tightly linked in species-rich grasslands. Philosophical Transactions of the Royal Society B: Biological Sciences 366:1346–1357.

Wilson, A. J., U. Gelin, M. C. Perron, and D. Réale. 2009. Indirect genetic effects and the evolution of aggression in a vertebrate system. Proceedings of the Royal Society B: Biological Sciences 276:533–541.

Wilson, A. J., D. Reale, M. N. Clements, M. M. Morrissey, E. Postma, C. A. Walling, L. E. Kruuk, et al. 2010. An ecologist’s guide to the animal model. Journal of animal ecology 79:13–26.

Wise, D. H. 2006. Cannibalism, Food Limitation, Intraspecific Competition, and the Regulation of Spider Populations. Annual Review of Entomology 51:441–465.

