## Supplementary materials for "Trait-specific indirect effects underlie variation in the response of spiders to cannibalistic social partners"

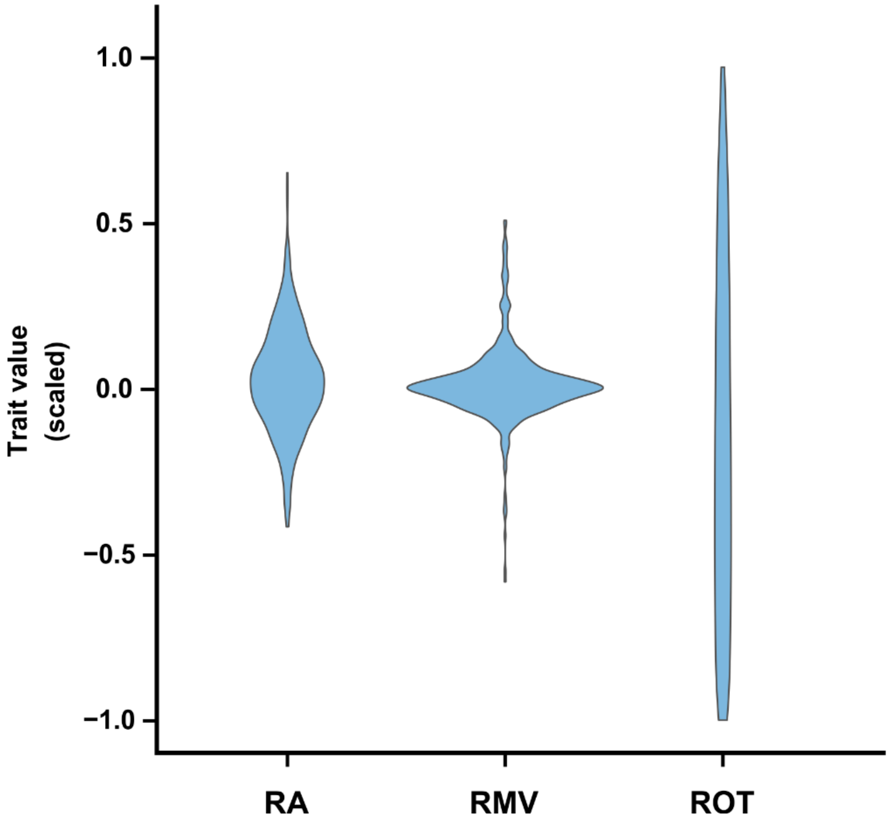


**Figure S1 – Violin plot showing the variation in trait values (standardized to mean zero and unit variance).** And its distribution, along the trait value range, in the three behavioral traits measured in response to conspecific cues, which constitute the composite behavior PC1. **RA** – relative activity; **RMV** – relative mean velocity; **ROT** – relative occupancy time.

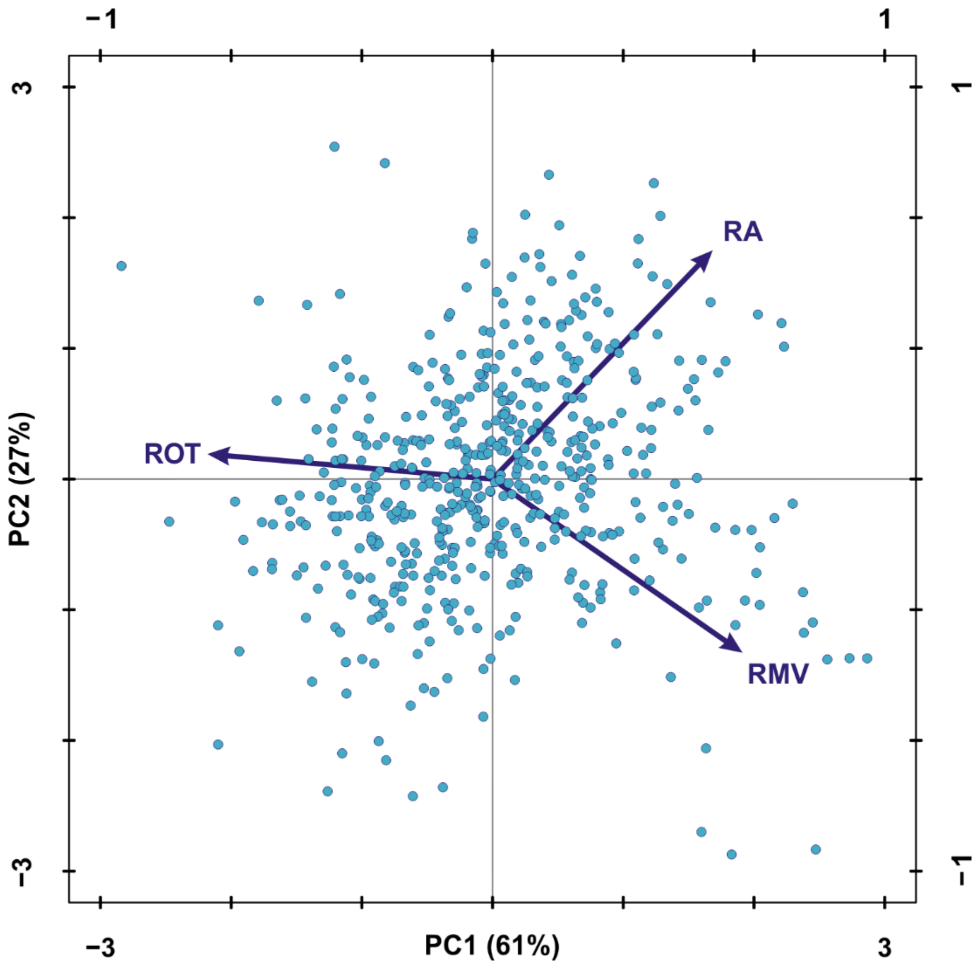


**Figure S2 – Principal component analysis on behavioral responses.** Explained variance: PC1, 61%; PC2, 27%. Each dot corresponds to one spider individual. **RA** – relative activity (loading on PC1, 0.66); **RMV** – relative mean velocity (loading on PC1, 0.77) and **ROT** – relative occupancy time (loading on PC1, -0.87).
